# Utilization of a cell-penetrating peptide-adaptor for delivery of HPV protein E2 into cervical cancer cells to arrest cell growth and promote cell death

**DOI:** 10.1101/2020.01.31.928358

**Authors:** Julia C. LeCher, Hope L. Didier, Robert L. Dickson, Lauren R. Slaughter, Juana C. Bejarano, Steven Ho, Scott J. Nowak, Carol A. Chrestensen, Jonathan L. McMurry

**Affiliations:** Department of Molecular & Cellular Biology, Kennesaw State University, Kennesaw, Georgia, United States of America; Department of Chemistry & Biochemistry, Kennesaw State University, Kennesaw, Georgia, United States of America

## Abstract

Cervical cancer is the second leading cause of cancer deaths in women worldwide. Human papillomavirus (HPV) is the causative agent of nearly all forms of cervical cancer, which arises upon viral integration into the host genome and concurrent loss of regulatory gene E2. E2 protein regulates viral oncoproteins E6 and E7. Loss of E2 upon viral integration results in unregulated expression and activity of E6 and E7, which promotes carcinogenesis. Previous studies using gene-based delivery show that reintroduction of E2 into cervical cancer cell lines can reduce proliferative capacity and promote apoptosis. However, owing in part to limitations on transfection *in vivo*, E2 reintroduction has yet to achieve therapeutic usefulness. A promising new approach is protein-based delivery systems utilizing cell-penetrating peptides (CPPs). CPPs readily traverse the plasma membrane and are able to carry with them biomolecular ‘cargos’ to which they are attached. Though more than two decades of research have been dedicated to their development for delivery of biomolecular therapeutics, the full potential of CPPs has yet to be realized as the field is hindered by the tendency of CPP-linked cargos to be trapped in endosomes as well as having significant off-target potential *in vivo*. Using a CPP-adaptor system that reversibly binds cargo thereby overcoming the endosomal entrapment that hampers other CPP methods, bioactive E2 protein was delivered into living cervical cancer cells, resulting in inhibition of cellular proliferation and promotion of cell death in a time- and dose-dependent manner. The results suggest that this nucleic acid- and virus-free delivery method could be harnessed to develop novel, effective protein therapeutics for treatment of cervical cancer.

## Introduction

Human papillomavirus is a sexually transmitted virus and the causative agent of multiple forms of cancer including cervical, vaginal, oropharyngeal, anal, penile and vulvar. Vaccines against the most common cancer-causing strains of HPV have been approved in the U.S. However, cervical cancer remains the second leading cause of cancer-related death in women worldwide [1]. Globally, this is partly attributed to a lack of access to preventive care and early detection, particularly in middle and low-income nations. Further, metastatic cervical cancer remains difficult to treat and retains high 5-year recurrence rates. In the U.S. alone, the 5-year survival rate for cervical cancer is 67%. A 92% survival rate has been achieved for early detection cases, but is only 17% in cases of late detection [2]. The most common means for treating metastatic cervical cancer are radiation and chemotherapy, but the past several years have seen a surge in clinical trials aimed at developing new immunotherapies to increase survival rates and reduce effective doses of traditional, harsher treatments. However, only one drug, Avastin, has been approved in the U.S. HPV-mediated cervical cancer thus remains a significant global burden and new treatment approaches are wanting.

A key event in over 80% of HPV-mediated cancers is viral integration into the human genome. During primary infection, HPV infects undifferentiated cells of the cervical basal epithelium. New virions exit from terminally differentiated cells in the outer layer of the cervical epithelium. The virus thus requires the proliferation and differentiation of host endodermal cells up the cervical epithelial wall for egress of new virions [3]. To insure this occurs, HPV encodes two proteins, E6 and E7, that inhibit apoptotic pathways and promote cellular proliferation, respectively [4, 5]. Another viral protein, E2, regulates E6 and E7 at the levels of both transcription and direct protein binding [6–8]. In over 80% of HPV carcinomas, the E2 open reading frame (ORF) serves as the primary site of viral integration, resulting in the loss of E2 but retention of E6 and E7 ORFs [9–13]. Subsequent unregulated overproduction of E6 and E7 leads to cellular changes promoting carcinogenesis. Loss of E2 is thus a critical event in the onset of many integrated HPV cancers.

Given its regulatory role of inhibiting E6 and E7, Hwang et al. hypothesized that replenishment of E2 in cervical cancer cells could prevent their proliferation and reverse their metastatic potential [14]. They and others demonstrated that reintroduction of E2 into cervical cancer cells could induce cell senescence [14–16]. Later work showed that E2 overexpression promotes apoptosis [17–21]. Currently there is no consensus in the field as to molecular events dictating one outcome over the other following E2 reintroduction. However promising, this approach has not become a viable treatment option for cancer patients, owing in great part to the need for gene delivery via transfection or viral vector delivery, a technical challenge in and of itself [22].

More recently, attempts have been made to overcome problems associated with gene delivery by developing protein replacement therapies whereby the protein product is directly delivered to cells or tissues in a controlled fashion. One of the most promising means for delivery of proteins into cells is cell-penetrating peptides (CPPs). CPPs are short peptides that can readily cross cell membranes and confer that ability on biomolecules to which they are attached [23–25]. CPP attachment is most commonly achieved via covalent bond or nonspecific hydrophobic interaction. However, these CPP-cargos often become trapped in endosomes upon cellular entry and, as a result, become targeted for degradation, resulting in cargo destruction rather than delivery to the cytoplasm or subcellular destination [26].

In 2004, Roeder et al described the use of the HIV CPP, VP22, to deliver VP22:E2 fusion proteins into cervical cancer cell lines for the induction of apoptosis [19]. In it and later studies, VP22:E2 fusion proteins were readily made from plasmids introduced into cells either via transfection or viral vector delivery [19–21]. Once translated, these fusion proteins were secreted from transformed cells and readily entered other neighboring cells to promote cell death [19–21].

In this study we describe an efficient, high-affinity reversible adaptor system for CPP-mediated delivery of E2 to cell interiors that exploits natural extra- and intracellular levels of calcium [27, 28]. The adaptor, TAT-CaM, consists of well-known CPP, TAT fused to a human calmodulin (CaM). Cargo proteins are engineered to contain a calmodulin binding sequence (CBS). TAT-CaM binds CBS-containing proteins with nM affinity in the presence of calcium and negligibly in its absence. Given that the extracellular environment contains relatively high levels of calcium, complexes remain tightly associated upon entry into the cell. However, during endosomal trafficking, calcium efflux results in cargo dissociation from TAT-CaM and subsequent release to the cytoplasm of living mammalian cells. Delivery is rapid, tunable and efficient. Further, at doses used in this study, TAT-CaM appears to be non-toxic, in agreement with a previous finding of measurable toxicity only at tens of μM and above [29].

Using the TAT-CaM adaptor system, the hypothesis that bioactive CBS-E2 protein delivered directly into living cervical cancer cells would inhibit cellular proliferation and/or promote cell death was tested. Following delivery, E2 showed distinct cell-cycle dependent subcellular localization patterns and was found in both the cytoplasm and the nucleus. In mitotic cells, E2 relocalized to regions of the cell associated with the mitotic spindle, a known biological activity [30]. As expected, E2 prohibited cellular proliferation and promoted cell death in a time and dose-dependent manner. The data support a model wherein E2 reduces cellular proliferation at low cell-to-peptide ratios and promotes cell death at high cell-to-peptide ratios. The data also further validate the TAT-CaM system and provide a new framework for delivery of E2 protein for the treatment of cervical carcinomas.

## Materials and Methods

### Generation and purification of CBS-E2 and TAT-CaM constructs

An *E. coli*-optimized synthetic gene encoding the E2 ORF from HPV-16 (GenBank: AAD33255.1) was cloned into pCAL-N-FLAG (Agilent Technologies, CA, USA), which contains a vector-encoded N-terminal calmodulin bind site (CBS). CBS-E2 and TAT-CaM were expressed and purified as previously described with slight modifications [27]. Briefly, CBS-E2 was expressed in ArcticExpress (DE3) *E. coli* cells (Agilent Technologies, USA) and purified by fast protein liquid chromatography using Calmodulin-Sepharose (GE Healthcare, USA). TAT-CaM was expressed in BL21(DE3)pLysS *E. coli* cells and purified to near-homogeneity by metal-affinity chromatography using TALON resin (Takara Bio, USA). After purification, proteins were dialyzed into calcium-containing binding buffer (10 mM HEPES, 150 mM NaCl, 2 mM CaCl_2_, 10% glycerol pH 7.4), sterilized via syringe-driven filtration through a 0.22 μm filter, flash frozen in liquid nitrogen and stored at −80°C until use.

### Biolayer Interferometry

Biolayer interferometry (BLI) experiments were performed on a FortéBio (Menlo Park, CA, USA) Octet QK essentially as previously described [27]. Biotinylated TAT-CaM was loaded onto streptavidin sensors for 300 s in binding buffer followed by a 180 s baseline measurement. Tethered TAT-CaM ligand was then exposed to analyte CBS-E2 and association was measured for 300 s. Two different dissociation phases followed, each 300 s in length. Ligand-analyte pairs were first exposed to binding buffer and were then challenged in binding buffer containing 10 mM ethylenediaminetetraacetic acid (EDTA). Baseline drift as measured by a parallel run in which a ligand-loaded sensor was exposed to buffer only was subtracted from each experimental run. Fast-on, slow-off binding was fit to a global 1:1 association-then-dissociation model and EDTA-induced rapid dissociation was separately fit to a one-phase exponential decay model using GraphPad Prism 5.02 software. Nonspecific binding as measured by a run of a sensor without ligand exposed to the highest concentration of CBS-E2 evinced negligible binding which was thus ignored in analysis.

### Cell culture

The HPV-16+ cell line SiHa (HTB-35) and the Human Microvascular Endothelial Cell line (HMEC; CRL-3243) were purchased from the American Type Culture Collection (ATCC). SiHa cells were cultured in glucose-free complete Dulbecco’s Minimal Eagle Media (DMEM; Gibco) with 10% fetal bovine serum (FBS; Atlas Biologicals) and 1 mM L-glutamine (Genclone). HMECs were cultured in MCDB131 media containing 10% FBS, 10 mM L-Glutamine, 10 ng/mL epidermal growth factor (EGF; Sigma), and 1 ug/mL Hydrocortisone (Sigma). Both cells lines were maintained in a humidified incubator at 37°C with 5% CO_2_ injection.

### Confocal microscopy

All confocal experiments were performed on an inverted Zeiss LSM700 confocal microscope equipped with a humidified incubator at 37°C with 5% CO_2_ injection essentially as previously described [27]. In short, SiHa cells were plated at ~50% confluency in 4-well Nunc Lab-Tek chambered coverglass wells (ThermoFisher) 16 hours prior to cell penetration assays. CBS-E2 cargos were labeled with DyLight 550 (ThermoFisher, USA), incubated with or without unlabeled TAT-CaM in equimolar amounts and brought to a total volume of 50 μL in binding buffer. Complexes were then added to 200 μL plain glucose-free media and introduced to cells at a final concentration of 1 uM. Uptake was performed in a humidified incubator at 37°C with 5% CO2 injection for 1 hour with periodic rocking (every 15 min) to ensure even distribution. After 1 hour, media were removed and cells were washed 5 times with calcium-containing phosphate buffered saline (PBS with 1mM CaCl_2_). Next, cells were counterstained with 2 μM CellTracker Green (5-chloromethylfluorescein diacetate dye) (ThermoFisher) followed by NucBlue (ThermoFisher) per manufacturer’s protocols to stain the cytoplasmic and nuclear compartments of the cells. After staining, cells were washed 3x with calcium-containing PBS and complete cell culture media was added to each well. Cells were imaged using a 40x EC Plan-Neofluar objective with a numerical aperture value of 1.3. Image analysis was performed as previously described [27].

### Analysis of cellular proliferation and cell death

Cellular proliferation was assayed by ability to metabolize 3-(4,5-dimethylthiazol-2-yl)-5-(3-carboxymethoxyphenyl)-2-(4-sulfophenyl)-2H-tetrazolium, inner salt (MTS) (Promega - CellTiter 96^®^ AQ_ueous_ One Solution Cell Proliferation Assay). In the same population of cells, cell death was assayed by release of LDH into cell culture media (Promega CytoTox 96^®^ Non-Radioactive Cytotoxicity Assay). Cells were seeded into 96-well plates at either 2.5 x10^3^ or 2.5 x10^4^ in 100 μL of phenol-red free cell culture media and allowed to adhere to the plate overnight (Day 0). The next day (Day 1) cells were treated with increasing amounts of CBS-E2 with equimolar TAT-CaM in binding buffer, TAT-CaM only (vehicle control), buffer (experimental control) or simply left untreated. After one hour treatments were removed and 100 μL of cell culture media were added to each well. Treatments were repeated at 24 and 48 hours. Every 24 hours, 50 μL of media were transferred to another 96 well plate and assayed for lactate dehydrogenase (LDH) per the manufacturer’s protocol. At 72 hours (Day 4) MTS reagent was added directly to cells and cells were assayed for MTS metabolism, also following the manufacturer’s protocol. A BioTek (BioTek Instruments, VT, USA) plate reader was used to measure OD_490_. Absorbance due to metabolic or LDH activity was calculated by subtracting background (cells with no reagent) from the total. Percent metabolic activity was then calculated using the following equation: (OD_treated_/OD_untreated_) x 100. The nature of this experimental approach (periodic sampling in living cells undergoing different rates of growth/death) required an alternative means to calculate LDH levels relative to controls. As such, LDH levels were represented by their respective OD_490_ values.

### Statistical Analyses

All analyses were performed using GraphPad Prism 8.0 software. Treated groups were compared to the untreated group using one-way ANOVA with Tukey’s correction for multiple comparisons. Deviation was calculated using standard error of the mean.

## Results

### CBS-E2 binds TAT-CaM with expected kinetics

Previous work validated that TAT-CaM binds model CBS-cargo proteins rapidly and stably in the presence of calcium and dissociates almost instantaneously and completely when calcium is removed [27, 28]. Given that relatively high calcium conditions prevail in the extracellular environment, that most mammalian cells maintain low intracellular calcium concentrations and that calcium and pH flux occurs during endosomal transport [31], the TAT-CaM methodology allows for rapid release of cargo from TAT-CaM once inside the cell, overcoming the ‘endosomal escape’ problem that has plagued the field [26]. In this study an E2 construct from HPV-16 that contained an N-terminal CBS tag was expressed and purified. Calcium-dependent binding kinetics of TAT-CaM with CBS-E2 were analyzed via biolayer interferometry (Fig 1). Fits to a single-state association-then-dissociation model (Fig 1A) yielded a calcium-replete K_D_ of 36 nM with k_on_ = 4.6 x 10^4^ M^-1^s^-1^ and k_off_ = 1.6 x 10^-3^ s^-1^. In the presence of the chelating agent EDTA, dissociation was very rapid (Fig 1B). k_off(EDTA)_ was 5.3 x 10^-2^ s^-1^. That the kinetic constants were highly similar to other cargos examined previously [27,28] validated the utility the approach for delivery of E2.

**Fig. 1.**
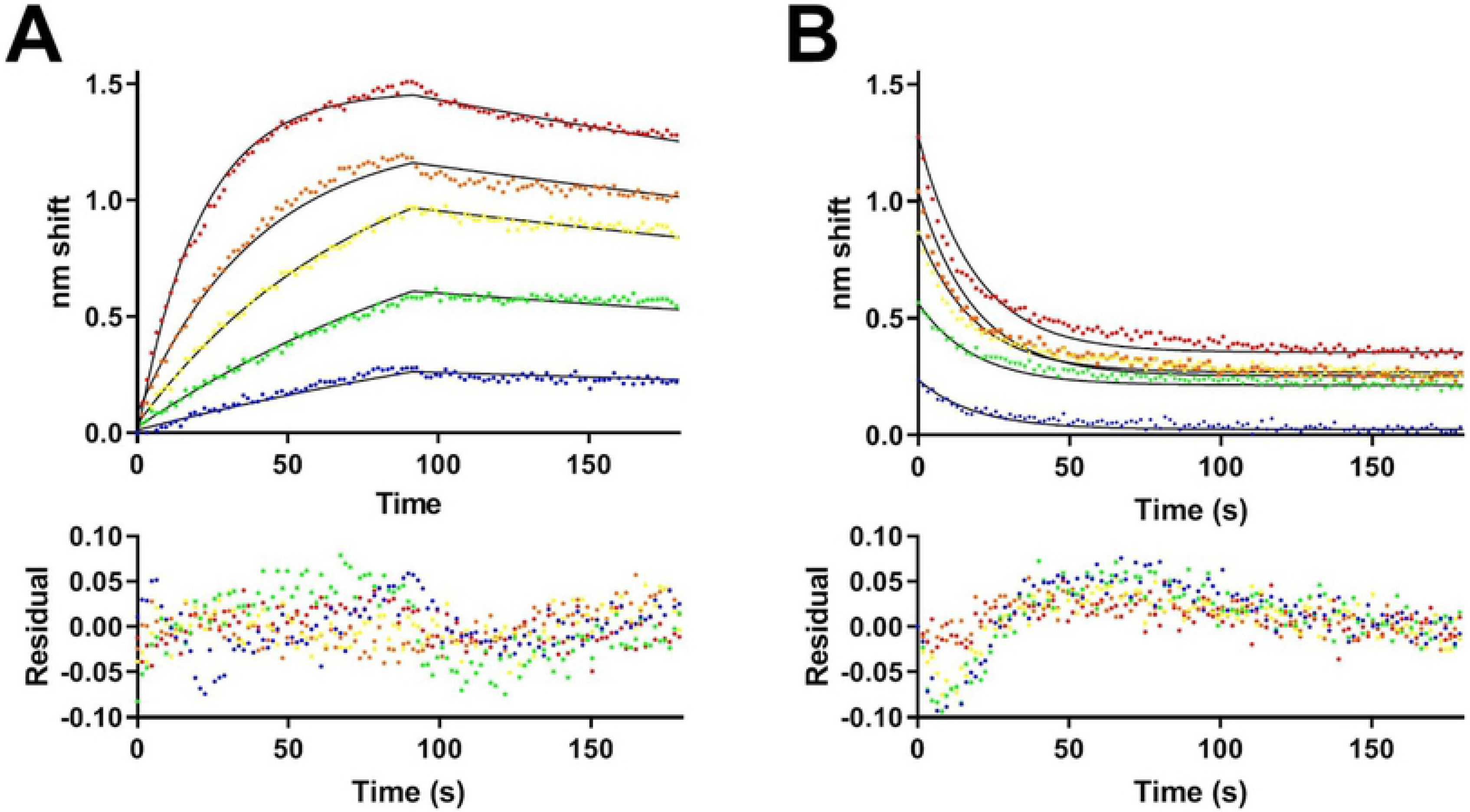
Biolayer interferometry analysis of TAT-CaM binding to CBS-E2. A, Association-then-dissociation experiment in which ligand TAT-CaM was exposed to varying concentrations of CBS-E2 prior to movement to buffer only at 90s (red, 1000 nM; orange, 500 nM; yellow, 250 nM, green, 125 nM, blue 63 nM). Data points are individual instrument readings. Lines represent best fits to a global single-state model. Residuals are shown below. B, The same samples after dissociation were moved to buffer containing 10 mM EDTA for monitoring of dissociation in the absence of Ca^2+^. Fits are to a global single-state exponential decay model.

### Live cell uptake and cellular redistribution of CBS-E2 post TAT-CaM-mediated delivery

TAT-CaM was used to deliver free bioactive E2 protein into the human HPV-16+ cervical cancer cell line SiHa. Given significant artifacts resulting from fixation that have confounded results in the past, live cell imaging in asynchronous populations of human cervical cancer cells was performed. *Z*-stacks were acquired via confocal microscopy and analyzed for intracellular delivery of fluorescently labeled CBS-E2 by TAT-CaM (Fig 2). To verify that TAT-CaM mediated entry, parallel control experiments without TAT-CaM were performed in which negligible signal from the fluorophore was observed (Fig 2A). In the presence of TAT-CaM, CBS-E2 was readily delivered into cells as evinced by fluorescence observed throughout the cell (Fig 2B).

**Fig. 2.**
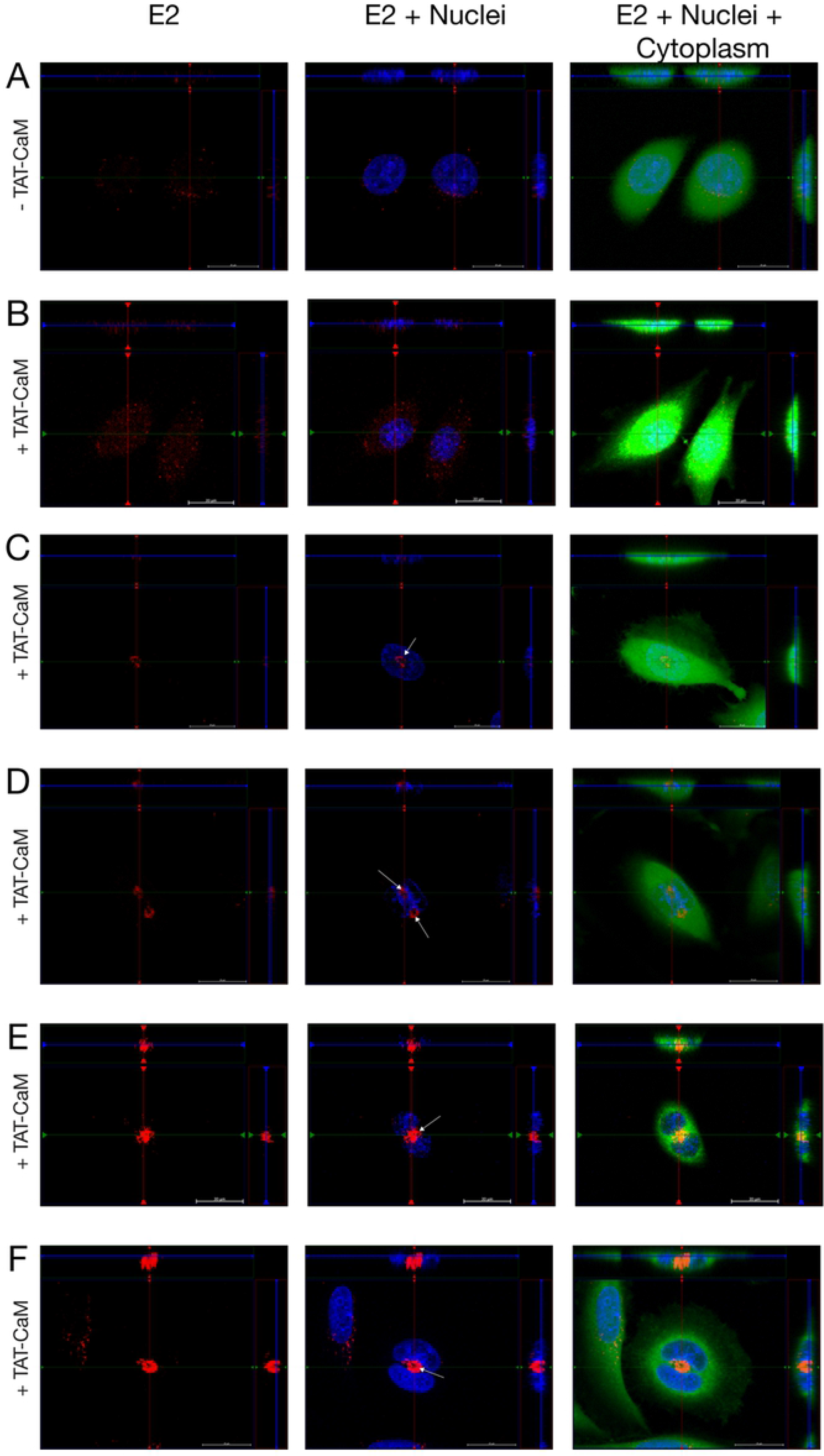
Delivery of CBS-E2 into living cervical cancer cells via the TAT-CaM adaptor. Cervical cancer cells were incubated with fluorescently labeled CBS-E2 cargo (red) in the absence (A) or presence (B-F) of equimolar TAT-CaM for 1 hr. Cells were counterstained with NucBlue (nuclei; blue) and CellTracker (cytoplasm; green). Images were generated on an inverted Zeiss LSM700 Confocal Microscope with Z-stack projections. Shown at the top and right of each image are orthogonal projections taken at the depth of the nucleus. A, B) Visualization of E2 in SiHa cells in asynchronous populations. C-F) Visualization of E2 in mitotically active cells. White arrows indicate redistribution and clustering of E2 to regions of the cell typically associated with the mitotic spindle apparatus.

One biological property of HPV E2 proteins is the ability to localize to the mitotic spindle during cellular division [30, 32]. While both low-risk (LR) and high-risk (HR) E2 proteins can associate with the spindle, the manner of association and resultant distribution pattern of LR and HR E2s during mitosis differ [30, 32]. High-risk HPV E2 proteins exhibit a dynamic relocalization pattern whereby they initially cluster at the asters at the onset of mitosis [30]. As the cell progresses through mitosis, E2 relocates to the midplane where it may associate with components of the Anaphase Promoting Complex (APC/C) [30, 32–34]. E2 remains at the midbody through cytokinesis. In concordance with these results, mitotic cells in cell-penetration experiments had a distinctive pattern of CBS-E2 redistribution to regions structurally similar to spindle fiber formations (white arrows; Fig 2C-F). Circular clusters of E2 formed on DNA were observed at the onset of mitosis (white arrows; Fig 2C, D), suggestive of localization to aster microtubules as previously described [30, 32]. In cells undergoing anaphase and telophase, CBS-E2 clustered on the midplane (white arrows; Fig 2E, F). These data demonstrate that CBS-E2 was delivered in bioactive form and that the CBS tag did not significantly affect activity.

### E2 inhibits cell progression and induces cell death in cervical cancer cells

Previous studies showed that transfection of cervical cancer cells with E2 is sufficient for induction of senescence or apoptosis within 3 days [14–18]. An experimental limitation to the use of gene delivery is lack of control of dose, i.e. how much protein is made in the cell after transfection or viral vector delivery. To test if CBS-E2 could induce senescence and/or cell death by direct protein delivery, experiments were designed to determine how much protein would be required over 3 days. As a starting point, 2.5 x 10^4^ cells were treated daily for 3 days with 1 or 4 μM doses of CBS-E2 and equimolar TAT-CaM. At 1 μM no significant effect on cells were observed after E2 delivery, while at 4 μM, there was a 28% reduction in metabolic activity on day 4 (Fig 3A). Total cell counts were also performed at day 4. Untreated and TAT-CaM only treated groups showed similar growth rates while cells treated with 4 μM E2 failed to proliferate (Fig 3B). Microscopic analysis of cells on day 4 further corroborated these findings (Fig 3C-E). Untreated and TAT-CaM treated cells exhibited normal morphology while E2 treated cells overwhelming became flattened out, with a loss in typical spindle-like morphology, and exhibited intracellular stress granule-like formations (Fig 3E). Collectively, these data support that 3 doses of E2 protein over a three-day time-period are sufficient to significantly reduce cellular proliferation within this population of cells.

**Fig. 3.**
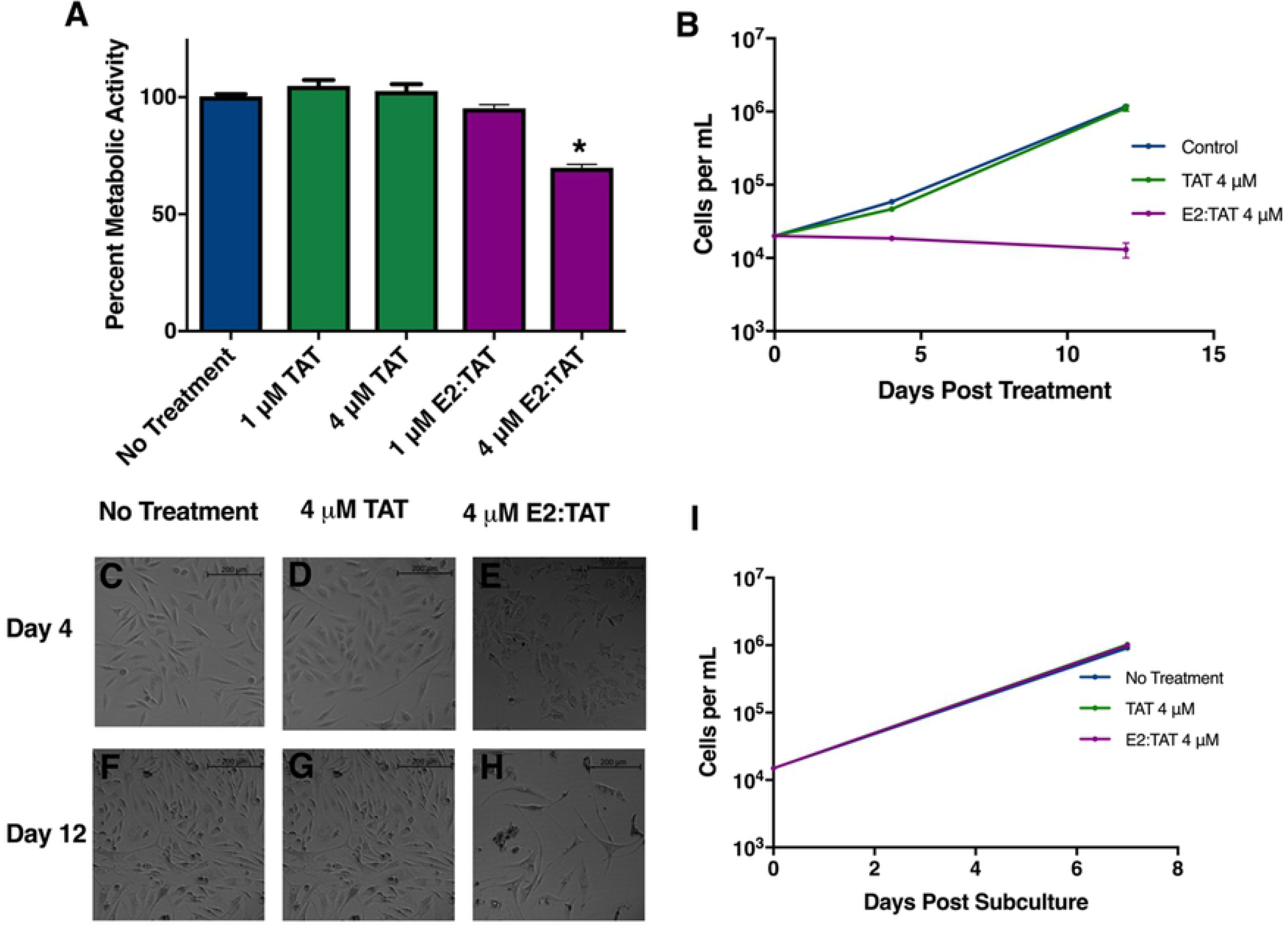
E2 delivery induces reversible growth arrest in cervical cancer cells. Cells were seeded at 2.5 x 10^4^ per well and treated once daily for three days with either 1 μM or 4 μM CBS-E2 in the presence of equimolar amounts of TAT-CaM. As a control, cells were either left untreated (negative control) or treated with TAT-CaM only (experimental control). A) MTS assay to assess cellular metabolic activity on day 4. A reduction in metabolic activity was tested for by One-way ANOVA with Dunnet’s correction for multiple comparisons *p= 0.03. n=9; shown SEM. B) On day 4 and day 12, cells were collected and counted on a hemocytometer. n=4. C-H) Micrographs of cells from each treatment group on day 4 and day 12 post treatment. Images are from the 4 μM treatments. I) On day 12 cells were collected and reseeded at equal density and cultured for an additional week after which they were collected and counted on a hemocytometer. n=4.

Persistence of phenotype was assayed by retaining cells in culture for an additional week with regular media changes. Over 12 days, with no additional E2 treatments, cells from the 4 μM treated group failed to proliferate while untreated cells and those dosed with TAT-CaM only retained normal doubling times (Fig 3B). Microscopic analysis supported these findings (Fig 3 F-H). In untreated and TAT-CaM treated groups, cells became over-confluent and crowded the wells (Fig 3F, G). E2-treated cells showed no visual evidence of doubling, however, some cells within the population regained normal spindle-like morphology (Fig 3H). We hypothesize that these cells might represent a subpopulation of harder to treat ‘persister’ cells. To test for this, cells were collected and re-seeded at equal density. After 7 days in culture, cells were collected and counted (Fig 3I). E2 treated cells regained normal growth kinetics (Fig 3I) and normal morphology (data not shown) indistinguishable from untreated or TAT-CaM treated cells. These data suggest that at the cell-to-peptide ratios employed, a sub-population of cells underwent senescence while others were seemingly unaffected or more resistant to E2’s effects.

To test the effect of cell-to-peptide ratios a dose-response assay was performed using the same protocol with the exception that the starting cell number was lowered 10-fold. While 0.1 μM doses had little effect, cells showed a dramatic reduction in metabolic activity at only 1 μM (75% loss; Fig 4A). Similar observations were made with 10 μM doses, suggesting that at doses >1 μM there is a ‘plateau effect,’ in that higher doses had no discernible effect (Fig 4A). Within the same population of cell, cell death was tested each day by measuring total LDH levels in the media. Results showed significantly high levels of LDH in all E2 treatment groups (Fig 4B). The much smaller level of LDH activity in controls was attributed to retention of normal growth rates leading to overconfluency. Next, E2’s ability to induce cell death in a non-cervical cancer human microvascular endothelial cell line (HMEC) was tested. The same LDH leakage assay was performed as above using the highest dose group of TAT:CaM & CBS-E2 (10 μM) in both SiHa and HMEC cell lines. SiHas showed significantly higher levels of cell death when compared with all other treatment groups while there was no discernable effect on HMECs following E2 delivery (Fig 5A). Microscopic analysis of cells on day 3 qualitatively corroborated these results (Fig 5 BI). Collectively, these data support the hypothesis that direct delivery of E2 protein into living cervical cancer cells can inhibit cellular proliferation or induce cell death and, further, suggest that these differential outcomes may be a function of dose. That E2 did not induce cell death in the HMEC cell line suggest a specificity for HPV+ cells.

**Fig. 4.**
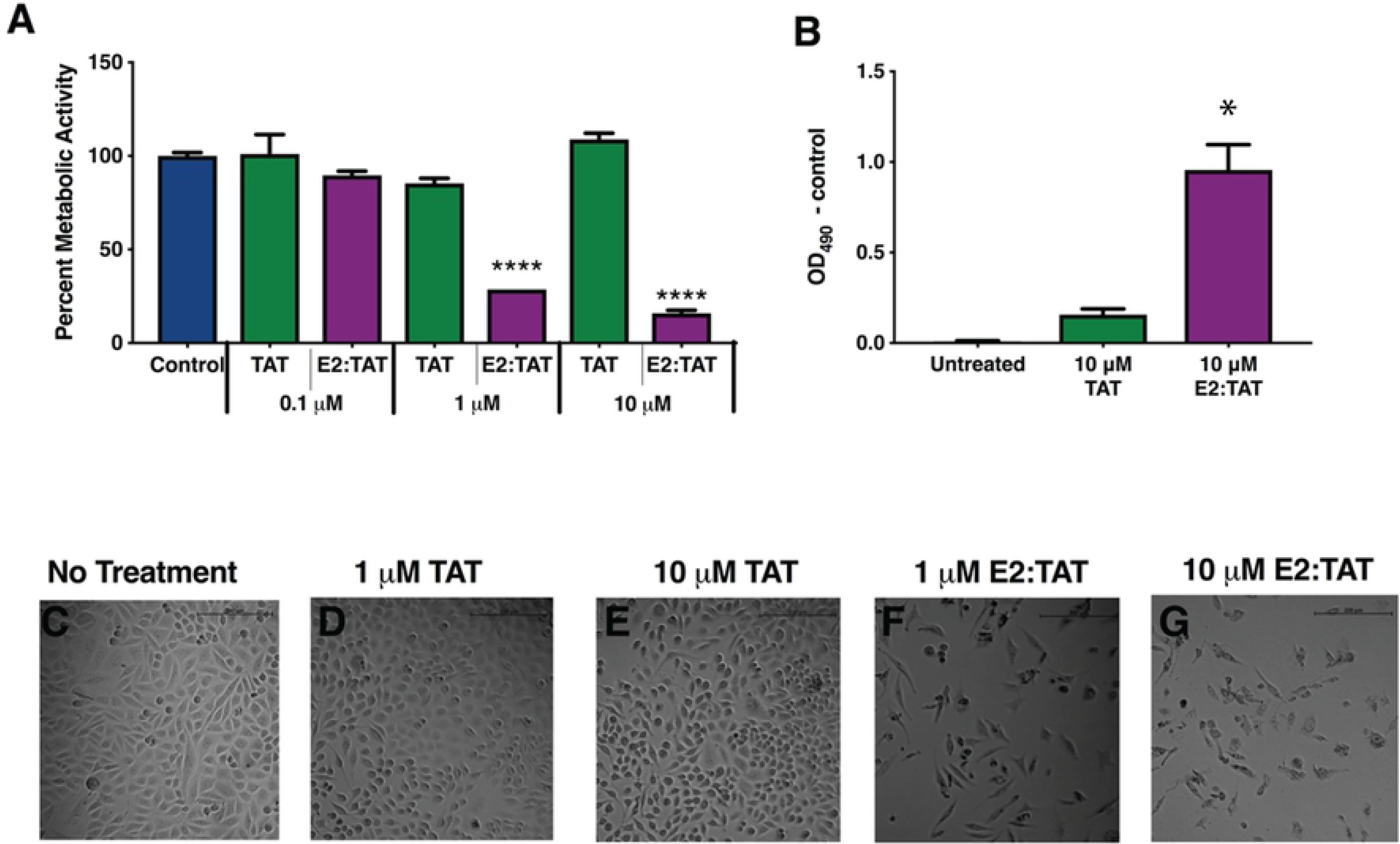
E2 induces cell death in cervical cancer cells. Cells were seeded at 2.5 x 10^3^ and treated once daily for three days with either 1, 3 or 10 μM E2 in the presence of equimolar amounts of TAT-CaM. As a control, cells were either left untreated (negative control) or treated with TAT-CaM only (experimental control). A) MTS assay to assess cellular metabolic activity on day 4. B) LDH leakage assay to assess cytotoxicity on day 4. A reduction in metabolic activity and percent cytotoxicity were tested for by One-way ANOVA with Dunnet’s correction for multiple comparisons. n=9; shown SEM. C-G) Micrographs of cells from each treatment group taken on day 4.

**Fig. 5.**
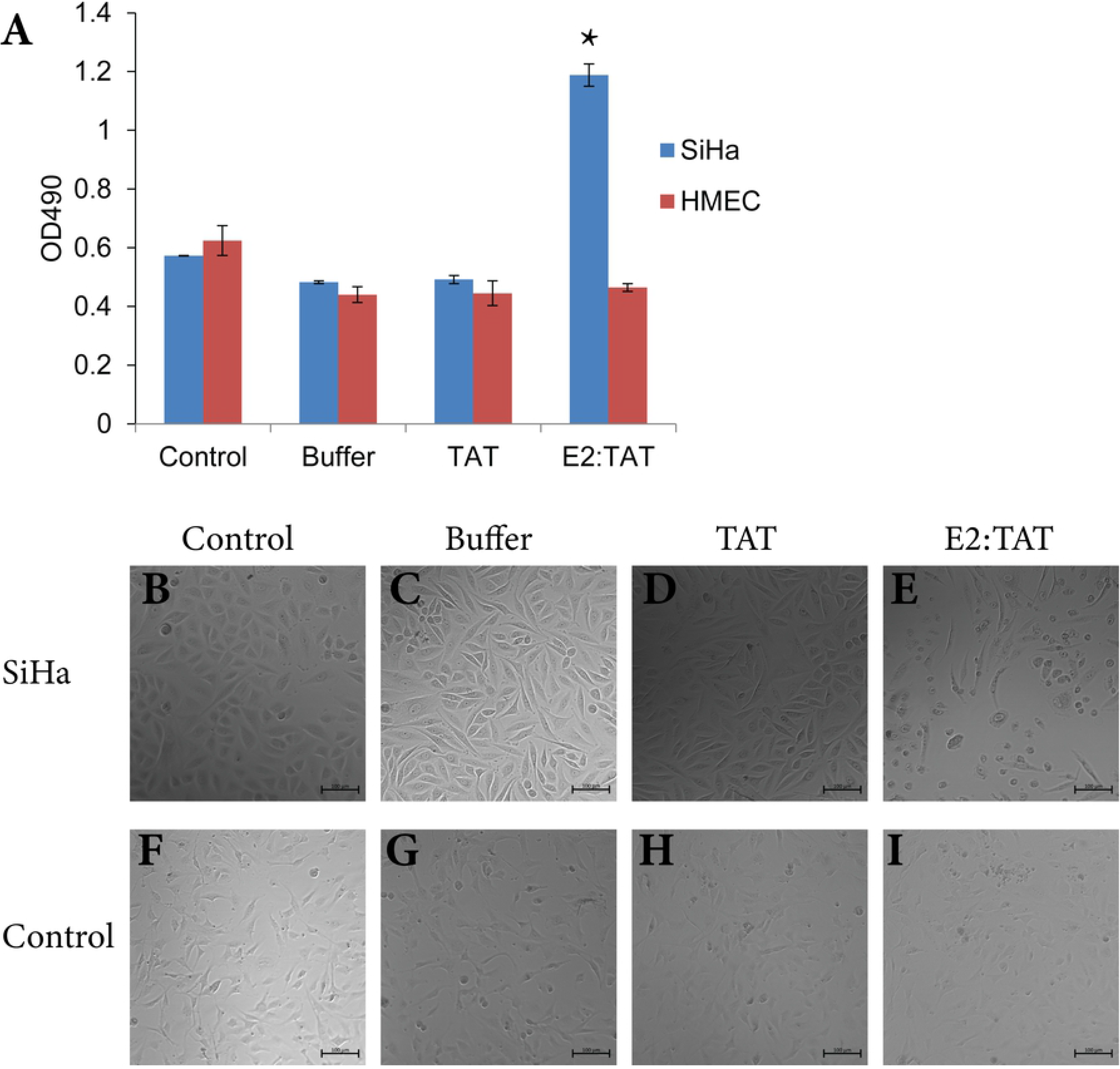
E2 does not induce cell death in Human Microvascular Endothelial Cells. SiHa and HMEC cells were seeded at 2.5 x 10^3^ and treated once daily for three days with 10 μM E2 in the presence of equimolar amounts of TAT-CaM. As a control, cells were either left untreated (negative control), treated with NEB buffer (treatment control), or treated with TAT-CaM only (experimental control). A) LDH leakage assay to assess cytotoxicity on day 4. A reduction in metabolic activity and percent cytotoxicity were tested for by One-way ANOVA with Dunnet’s correction for multiple comparisons. n=4; shown SD. B-I) Micrographs of cells from each treatment group taken on day 4.

## Discussion

This work sought to address the hypothesis that bioactive viral E2 protein could be delivered directly into living cervical cancer cells via the TAT-CaM CPP-adaptor system to inhibit cellular proliferation and/or promote cell death. CBS-E2 showed the expected high affinity, calcium-dependent binding kinetics with TAT-CaM (Fig 1), well within the range of constants previously determined for TAT-CaM other cargo proteins [27, 28]. Plateaus observed in the EDTA dissociation phase are confounding in that complete dissociation ought result in a plateau of 0 nm shift given that non-specific binding of analyte to the sensor was near zero. However, similar plateaus have been previously observed with CaM and analytes in BLI and were attributed to partial denaturation of the proteins, perhaps a result of tethering to the sensor [35]. Another contributor is uncertainty of the value of Y_0_ (Y at the beginning of dissociation) as dissociation is very rapid and the instrument takes a reading only every 1.6 s. For very fast processes, there is also often a discontinuity between the end of one phase, in this case dissociation in Ca^2+^, and the next, dissociation in EDTA. Indeed, the residuals (Fig 1B) indicate the poorest fit at the outset of dissociation, though they remain within the range of normal for the BLI instrument. Regardless of the idiosyncratic uncertainties inherent in the measurements, the kinetics of the interaction were as expected and suitable for delivery of CBS-E2 into cells.

TAT-CaM readily delivered CBS-E2 constructs into the HPV-16+ cervical cancer cell line SiHa (Fig. 2). In the course of the cell penetration assays a distinctive and repeatable pattern of intracellular relocalization in mitotic cells to regions associated with mitotic spindle fibers was observed (Fig. 2C-F). The pattern of spindle fiber association was congruent was previously published reports for high-risk E2 proteins in HPV+ cell lines [30, 32]. These data suggest that CBS-E2 protein constructs are biological active post TAT-CaM mediated intracellular delivery.

Previous E2 reintroduction studies in HPV16+ & 18+ cell lines using transfection or viral vector delivery showed senescence and/or cell death [14–18]. Following introduction of CBS-E2 protein into SiHa cells, both phenotypes were observed. The conditions dictating one outcome over the other within this study were attributed to cell-to-peptide ratios; at low ratios senescence may be favored while at high ratios cell death is (Fig. 3 and 4). E2’s effects are specific to HPV+ cervical cancer cells as E2 delivery into the non-cancerous non-cervical HMEC cell line failed to induce cell death (Fig 5), which is not surprising considering that the mode of action for E2 is inhibition of other viral proteins that are not present in HMEC. In short, the data presented herein support the hypothesis and provide a new framework to study the biological activity and therapeutic potential of HPV viral protein E2, though the mechanism(s) underlying these differential outcomes is still unknown. Utilizing viral vector delivery of E2, Desaintes, et al. revealed that both senescence and apoptosis could occur within the same cellular population and postulated that these outcomes may be the result of the amount of E2 being made within the cell [17]. The results of the present study, i.e. cell death was not detected at the low cell-to-peptide ratio, suggest a dose-dependent effect whereby E2 inhibits cellular proliferation at low ratios and promotes cell death at high ratios. A possible explanation for the differential effects of E2 is persistence of resistant cells within the heterogeneous SiHa population. Correlation between cell subtype and E2 effects may further illuminate E2’s efficacy both at different stages in carcinogenesis as well as within a heterogeneous tumor population. It is hoped that CPP-adaptor-mediated delivery can be used to address these questions in a future study.

A final consideration is the manner of E2-mediated cell death. Overwhelmingly, cell death after E6 and/or E7 knockdown either via E2 reintroduction, mRNA silencing or gene editing has been attributed to apoptosis [16, 38, 40–52]. In 2017 Shankar et al. reported a more robust analysis of cell death following TALEN-mediated cleavage of E7 and showed that cell death was attributable to a necrosis-like phenotype [53]. While seemingly in contrast to a wealth of other studies, previous work relied predominantly on the use of annexin V and propidium iodide staining to investigate the manner of cell death. Though these techniques are well vetted for the detection of apoptotic cells they are unable to exclude necrosis-like cell death. Thus, it is feasible that alternative forms of cell death were overlooked in some instances. In the current study, LDH leakage from cells, which also fails to discriminate between different forms of cell death, was assayed. As an indicator of apoptosis, cleavage of caspases 8 and 3 were assayed (data not shown). While caspase cleavage was not readily detectable, we were unable to rule out apoptosis as the manner of cell death. Currently, experiments are underway to parse out manner of cell death after E2 protein delivery. As more sensitive and robust assays of cell death are employed, both apoptosis and necrosis occurring after E2 delivery may be observed.

The manner of cell death is a critical consideration in the design of therapeutics for the treatment of carcinomas. Tumors are known to evade the immune response by preventing the release of immunostimulants and down-regulating surface antigens [54]. Induction of necrosis in carcinomas is an attractive approach as it is a form of immunogenic cell death (ICD) that can promote the release of immunomodulatory molecules. In concert with appropriate immunotherapies, this approach may stimulate immunodetection and promote long-term immunosurveillance of tumors. Indeed, great strides have been made employing this combinatorial approach in preclinical studies employing oncolytic viruses [55].

In conclusion, this work lays the foundation for a new approach towards our understanding of the biology of HPV-mediated cervical cancer and studying the specific and interrelated roles of viral proteins in proliferation, senescence and cell death.

## Acknowledgements

We are grateful to Dr. Martin L. Hudson for helpful discussions and assistance in rendering Fig 2.

